# Multi-generational silencing dynamics control cell aging

**DOI:** 10.1101/102210

**Authors:** Yang Li, Meng Jin, Richard O’Laughlin, Philip Bittihn, Lev S. Tsimring, Lorraine Pillus, Jeff Hasty, Nan Hao

## Abstract

Cellular aging plays an important role in many diseases, such as cancers, metabolic syndromes and neurodegenerative disorders. There has been steady progress in identifying aging-related factors such as reactive oxygen species and genomic instability, yet an emerging challenge is to reconcile the contributions of these factors with the fact that genetically identical cells can age at significantly different rates. Such complexity requires single-cell analyses designed to unravel the interplay of aging dynamics and cell-to-cell variability. Here we use novel microfluidic technologies to track the replicative aging of single yeast cells and reveal that the temporal patterns of heterochromatin silencing loss regulate cellular lifespan. We found that cells show sporadic waves of silencing loss in the heterochromatic ribosomal DNA (rDNA) during the early phases of aging, followed by sustained loss of silencing preceding cell death. Isogenic cells have different lengths of the early intermittent silencing phase that largely determine their final lifespans. Combining computational modeling and experimental approaches, we found that the intermittent silencing dynamics is important for longevity and is dependent on the conserved Sir2 deacetylase, whereas either sustained silencing or sustained loss of silencing shortens lifespan. These findings reveal, for the first time, that the temporal patterns of a key molecular process can directly influence cellular aging and thus could provide guidance for the design of temporally controlled strategies to extend lifespan.

**Significance:** Aging is an inevitable consequence of living, and with it comes increased morbidity and mortality. Novel approaches to mitigating age-related chronic diseases demand a better understanding of the biology of aging. Studies in model organisms have identified many conserved molecular factors that influence aging. The emerging challenge is to understand how these factors interact and change dynamically to drive aging. Using multidisciplinary technologies, we have revealed a sirtuin-dependent intermittent pattern of chromatin silencing during yeast aging that is crucial for longevity. Our findings highlight the important role of silencing dynamics in aging, which deserves careful consideration when designing schemes to delay or reverse aging by modulating sirtuins and silencing.

## Main Text

Cellular aging is generally driven by the accumulation of genetic and cellular damage (1, 2). Although much progress has been made in identifying molecular factors that influence lifespan, what remains sorely missing is an understanding of how these factors interact and change dynamically during the aging process. This is in part due to the fact that aging is a complex process wherein isogenic cells have various intrinsic causes of aging and widely different rates of aging. As a result, static population-based approaches could be insufficient to fully reveal sophisticated dynamic changes during aging. Recent developments in single-cell analyses to unravel the interplay of cellular dynamics and variability hold the promise to answer that challenge (3–5). Here we chose the replicative aging of yeast *S.cerevisiae* as a model and exploited novel quantitative biology technologies to study the dynamics of molecular processes that control aging at the single-cell level.

## Results

Replicative aging of yeast is measured as the number of daughter cells produced before the death of a mother cell (6). The conventional method for studying yeast aging requires laborious manual separation of daughter cells from mother cells after each division and does not allow tracking of molecular processes over multiple generations during aging (7). Recent advances in microfluidics technology have automated cell separation and enabled continuous single-cell measurements during aging (8–14). Building upon these efforts, we developed a new microfluidic aging device. The device traps mother cells at the bottom of finger-shaped chambers, allowing them to bud continuously, while daughter cells are removed via a waste port. Each chamber also has a small opening at the bottom, allowing daughter removal when mother cells switch budding direction (Fig. 1A, B and Movie S1). The long trapping chambers allow tracking of each daughter cell during its first several divisions, useful for monitoring age-related daughter morphologies. Furthermore, dynamic experiments involving precise step changes in media conditions can be conducted using this device. In validating the device, we confirmed that the majority of loaded cells are exponentially-growing newborn or young cells and the Replicative Lifespans (RLS) measured using the device are comparable to those from classical microdissection (15, 16) (SI Appendix, Fig. S1A-D).

**Figure 1.**
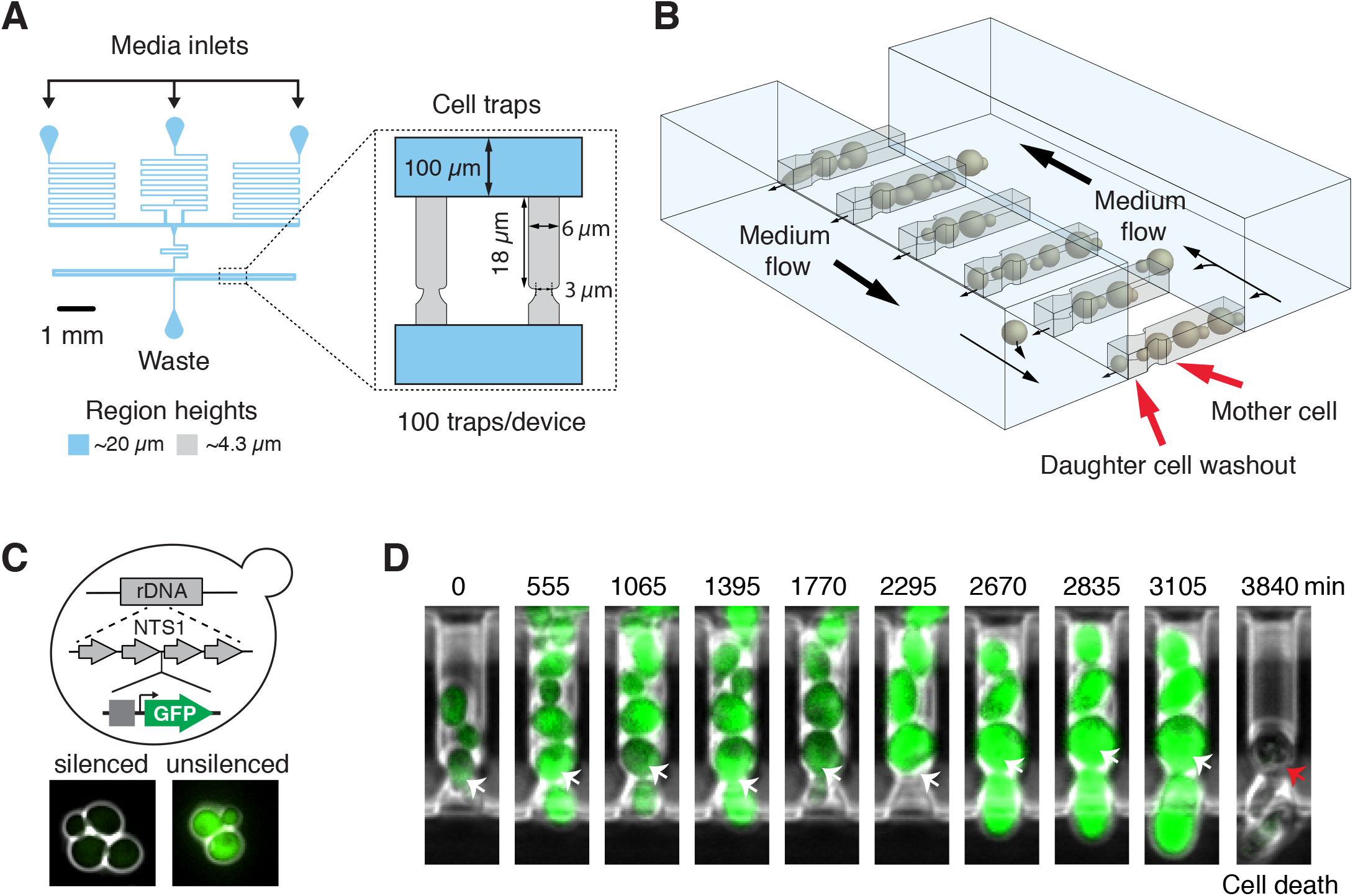
Tracking the dynamics of chromatin silencing in single aging cells using a new microfluidic device. (A) The design layout of the device. “Media” points to media inlets; “Cell trapping area” contains cell traps for single mother cells. (B) A schematic of multiple, parallel cell traps. Mother cells stay at the bottom of finger-shaped chambers while daughters are pushed out. Details of the device are provided in SI Appendix. (C) Illustrative schematic of the reporter for chromatin silencing. A GFP reporter under a strong constitutive promoter *(TDH3)* is inserted at an NTS1 repeat within the rDNA. Chromatin silencing at the rDNA is reflected by fluorescence intensity: low – silenced, high – decreased silencing. Details of the reporter are provided in SI Appendix. (D) Time-lapse images of a single cell trap throughout an entire lifespan. Arrows point to the mother cell.

Genome instability has been considered a significant causal factor of cell aging (1, 17). A major contributor to the maintenance of genome stability is chromatin silencing, which causes a locally inaccessible heterochromatin structure that represses transcription, recombination and DNA damage. The heterochromatic regions in yeast include the silent mating-type (HM) loci, rDNA and subtelomeric regions (18). Among them, the rDNA region on chromosome XII consists of 100~200 tandem repeats and is a particularly fragile genomic site, the stability of which closely connects to the RLS (19–21). Previous studies showed that silencing loss induced by chemical or genetic perturbations leads to increased recombination at the rDNA region and shorter lifespans (22, 23). However, the dynamic changes of rDNA silencing during cell aging remain largely unknown. To measure silencing in single aging cells, we constructed a strain with a fluorescent reporter gene under the control of a constitutive *TDH3* promoter at a nontranscribed spacer region (NTS1) of rDNA. Because expression of the reporter gene is repressed by silencing, decreased fluorescence indicates enhanced silencing, whereas increased fluorescence indicates reduced silencing (24, 25) (Fig. 1C). We observed that cells with the rDNA reporter gene exhibit weak fluorescence; in contrast, cells carrying the same reporter at the *URA3* locus, which is not subject to silencing, show very high fluorescence. In addition, deletion of *SIR2*, which is required for rDNA silencing (18), yields significantly increased reporter expression at the rDNA, but not at *URA3* (SI Appendix, Fig. S1E).

Using the microfluidic device and the reporter, we tracked the dynamics of rDNA silencing throughout the entire lifespans of individual cells (Fig. 1D; Fig. 2) We found intermittent fluorescence increases in most cells, indicating sporadic silencing loss during aging. About half (~46%) of the cells, during later stages of aging, continuously produced daughter cells with a characteristic elongated morphology until death (Fig. 2A, blue arrows). These cells also exhibited a sustained and dramatic increase in fluorescence, indicating sustained loss of silencing in aged cells (Fig. 2A, color map). In contrast, the other half of the cells, at later phases of their lifespans, continuously produced small round daughter cells (Fig. 2B, blue arrows) with sharply increased cell cycle length. These cells had a shorter average lifespan than the other aging type (with a mean RLS of 18, compared to 24) and did not show sustained silencing loss during aging (Fig. 2B, color map). These two distinct types of age-associated phenotypic changes suggest different molecular causes of aging in isogenic cells (8, 9, 14). Previous studies showed that the aging phenotype with small round daughters could be related to an age-dependent mitochondrial dysfunction (9, 12), but the molecular mechanisms underlying the other aging type characterized by elongated daughters remain largely unclear. Our results revealed that the sustained rDNA silencing loss, which can lead to genome instability (18), is specifically associated with the aging phenotype featured by elongated daughters. In support of this, young mother cells can also sporadically produce a few elongated daughters, the occurrence of which correlates with the transient silencing loss during the early phases of their lifespans (Fig. 2C). In addition, cross-correlation analysis revealed a ~140 min time delay between the occurrence of silencing loss in mother cells and the production of elongated daughters (SI Appendix, Fig. S2). This temporal order suggested a potential causal relationship between silencing loss and the elongated daughter phenotype. In this work, we focused our analysis on the dynamics and heterogeneity of the type of aging process with sustained silencing loss and elongated daughters.

**Figure 2.**
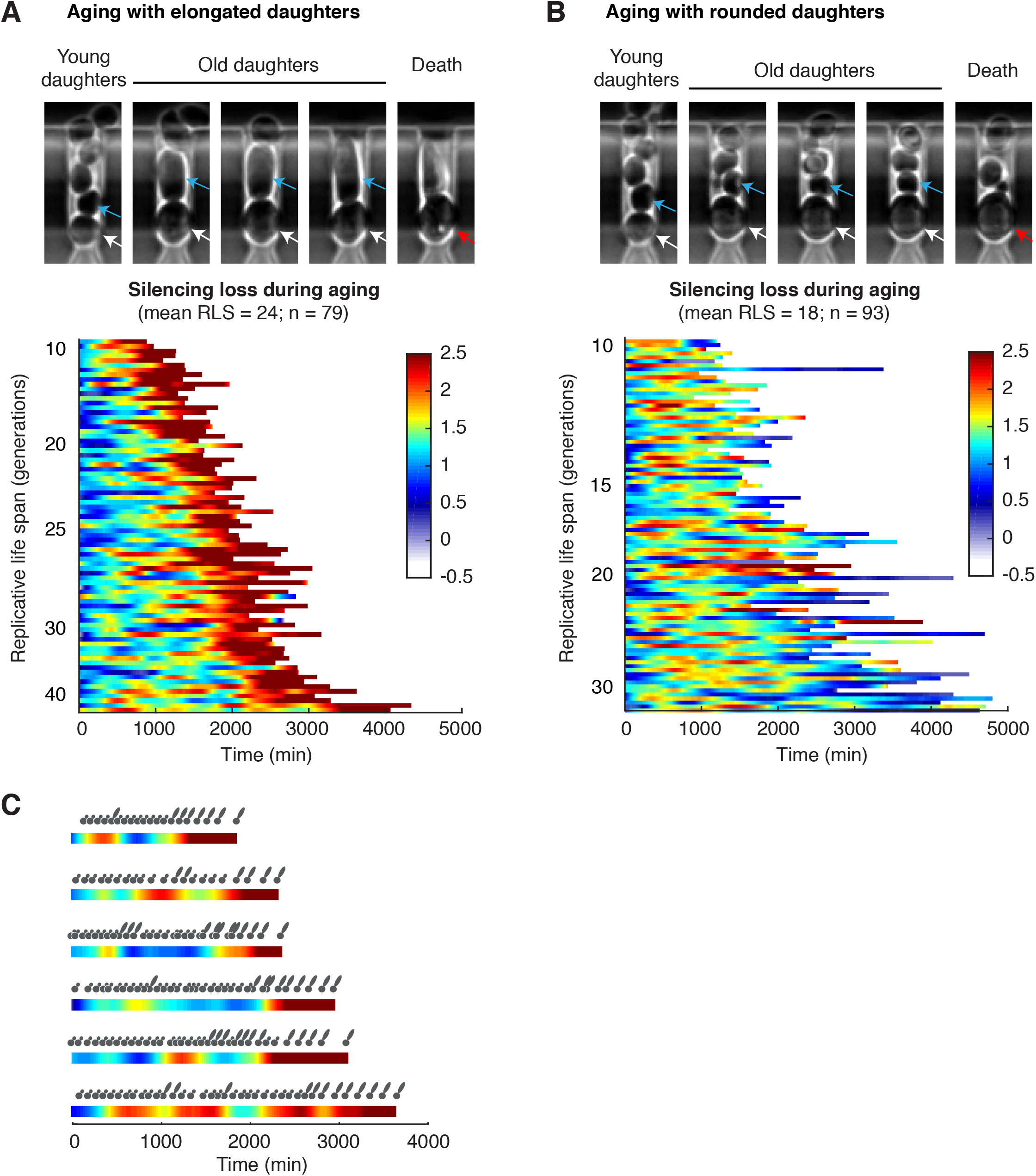
Dynamic patterns of silencing loss during aging. (A) Dynamics of silencing loss in cells aging with elongated daughters. Top: representative images of cell aging and death with elongated daughters. Blue arrows point to daughter cells; white arrows point to the living mother cell; red arrow points to the dead mother cell. Bottom: single-cell color map trajectories of reporter fluorescence. Each row represents the time trace of a single cell throughout its lifespan. Color represents the normalized fluorescence intensity as indicated in the color bar, which is used throughout to allow comparisons between different conditions. Cells are sorted based on their RLSs. (B) Dynamics of silencing loss in cells aging with rounded daughters. (C) Daughter cell morphology is correlated with the silencing state of mother cells. Color map trajectories of representative mothers are aligned with the morphology (round vs elongated) of daughters produced throughout their lifespans. A cross-correlation analysis of daughter morphology and mother silencing dynamics is shown in SI Appendix, Fig. S2.

To exclude the possibility that the observed fluorescence patterns are caused by age-associated global effects on gene expression (26), we simultaneously monitored two distinguishable fluorescent reporters inserted at the rDNA and at *URA3*. Whereas the rDNA reporter showed early sporadic and late sustained induction of fluorescence, the reporter at *URA3* exhibited relatively constant fluorescence during aging (SI Appendix, Fig. S3A). To confirm that the observed silencing dynamics are not specific to the NTS1 region, we measured the reporter response at another non-transcribed spacer region of rDNA (NTS2) and observed similar dynamic patterns as those found at NTS1 (SI Appendix, Fig. S3B). Together, these results validate the reporter responses during aging. We observed very rare events of recombination or extrachromosomal rDNA circles of the reporter gene, which can be easily distinguished from silencing loss in single-cell time traces (SI Appendix, Fig. S3C). We have excluded those cells from the analysis.

To evaluate how dynamic patterns of chromatin silencing influence cell aging, we quantitatively analyzed the time traces of silencing loss in individual cells producing elongated daughters before death. With very diverse lifespans ranging from 9 to 48 generations, all of the cells show sustained silencing loss toward later stages of aging. Most cells also exhibit early sporadic waves of silencing loss, each of which spans multiple cell divisions (Fig. 3A; Movie S2). This unprecedented long-wavelength dynamics is distinct from most previously characterized molecular pulses, which are on timescales faster than or close to a cell cycle (5). We further dissected each single-cell time trace into two phases: an early phase with sporadic silencing loss and a late phase with sustained silencing loss (Fig 3B, “Intermittent Phase” and “Sustained Phase”). The length of Intermittent Phase (or the number of silencing waves) is highly variable among cells (Coefficient of Variation – CV: 0.63; SI Appendix, Fig. S4A) and correlates closely with final lifespan, suggesting that the longevity of a cell is largely determined by the time it stays in this phase (Fig. 3C, left). Long-lived cells generally have a longer Intermittent Phase and produce more silencing waves than short-lived cells (Fig. 3A; SI Appendix, Fig. S4B and D). In contrast, the length of Sustained Phase is more uniform among cells (CV: 0.29; SI Appendix, Fig. S4A) and shows little relationship with lifespan, suggesting that sustained silencing loss defines cell death within a relatively constant period of time (Fig. 3C, right). We further quantified the rise time of each fluorescence increase in single aging cells (the duration of silencing loss; Fig. 3B, t_1_ and t_2_) and found a significant difference between the durations of early sporadic and later sustained silencing loss: a sporadic silencing loss on average lasts for ~ 300 minutes, whereas sustained silencing loss lasts for ~ 1200 minutes until death (Fig. 3D). Moreover, sporadic waves of silencing loss show modest effects on the basal silencing level during aging, as indicated by the trough levels of silencing loss pulses in single-cell time traces (SI Appendix, Fig. S4B and C), yet do not contribute additively onto sustained silencing loss to inducing cell death (SI Appendix, Fig. S4E).

**Figure 3.**
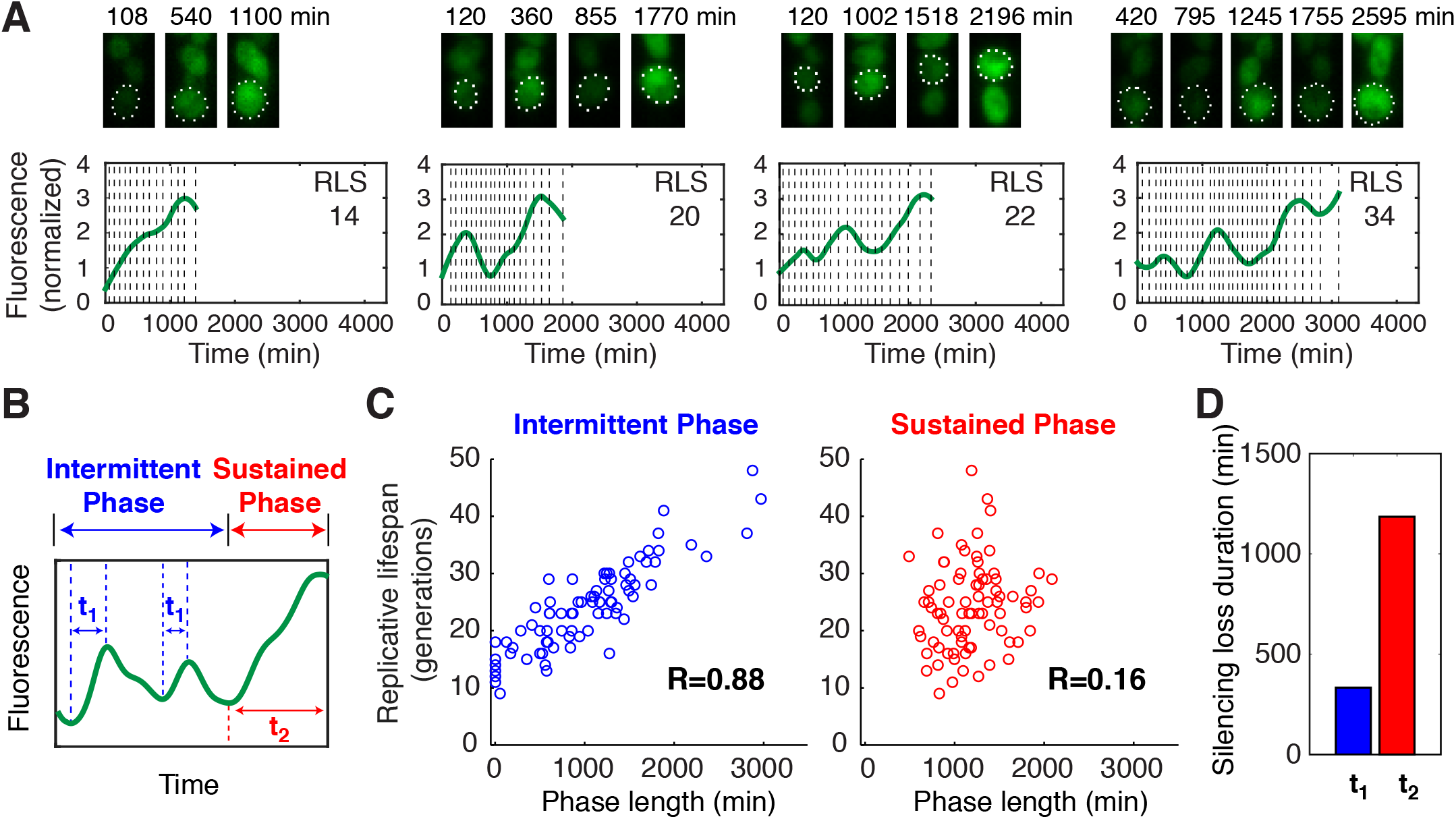
A quantitative analysis of silencing loss dynamics in single cells. (A) Dynamics of silencing loss in representative single cells with different lifespans. For each cell, (top) time-lapse images for the cell have been shown with (bottom) the fluorescence time trace throughout its lifespan. Vertical dashed line represents each division time of the cell, in which the distance between two adjacent dashed lines indicates the cell cycle length. Reporter fluorescence is normalized to the baseline level. (B) A schematic illustrates the dissection of two phases based on the silencing loss dynamics. The rise times of early sporadic and final sustained silencing loss are defined as “t1” and “t2”, respectively. (C) Scatter plots showing the relationships between the length of (left) Intermittent Phase or (right) Sustained Phase and lifespan at the single-cell level. Single-cell data are from Fig. 2A. Each circle represents a single cell. Correlation coefficient (R) is calculated and shown. (D) Bar graph showing the average durations of (blue) early sporadic and (red) final sustained silencing loss.

Together, our analyses reveal that cells undergo spontaneous silencing loss during aging. The early phase of aging features a reversible process, in which cells can effectively re-establish silencing and produce non-detrimental short waves of silencing loss. The late phase is irreversible: aged cells cannot re-establish silencing (27), resulting in sustained silencing loss and death. Individual cells may have different intrinsic capacities to maintain the reversible phase and thereby the ultimate lifespan.

To provide a quantitative framework for understanding aging dynamics, we developed a simple phenomenological model. The model postulates that an aging cell can be in one of two states: state 0 is the silenced state in which it produces normal daughters, and state 1 is the silencing loss state with elongated daughters (Fig. 4A). The transitions between the states are characterized by probabilities *p*_01_ and *p*_10_ that depend linearly on the cell age (Fig. 4B). We also assume that in the silencing loss state (state 1), a damage factor *D* accumulates uniformly and the probability of cell death is proportional to *D* and therefore to the number of generations a cell spends continuously in state 1 (Fig. 4C). In the silenced state (state 0), *D* is set to zero. We fit the model only using the experimental data on phenotypic changes and simulated this model stochastically. The model reproduced the main statistical properties of age-dependent phenotypic changes and RLS remarkably well (Fig. 4C-F). We also generated individual cell state trajectories (Fig. 4G) that qualitatively and quantitatively reproduce the data in Fig. 2A. To predict how silencing dynamics influence aging, we further simulated the effects of an induced silencing loss. Whereas a short pulse of silencing loss does not affect lifespan, a sustained silencing loss dramatically shortens lifespan (Fig. 4H and I). To test these predictions, we set out to modify silencing dynamics using genetic or chemical perturbations.

**Figure 4.**
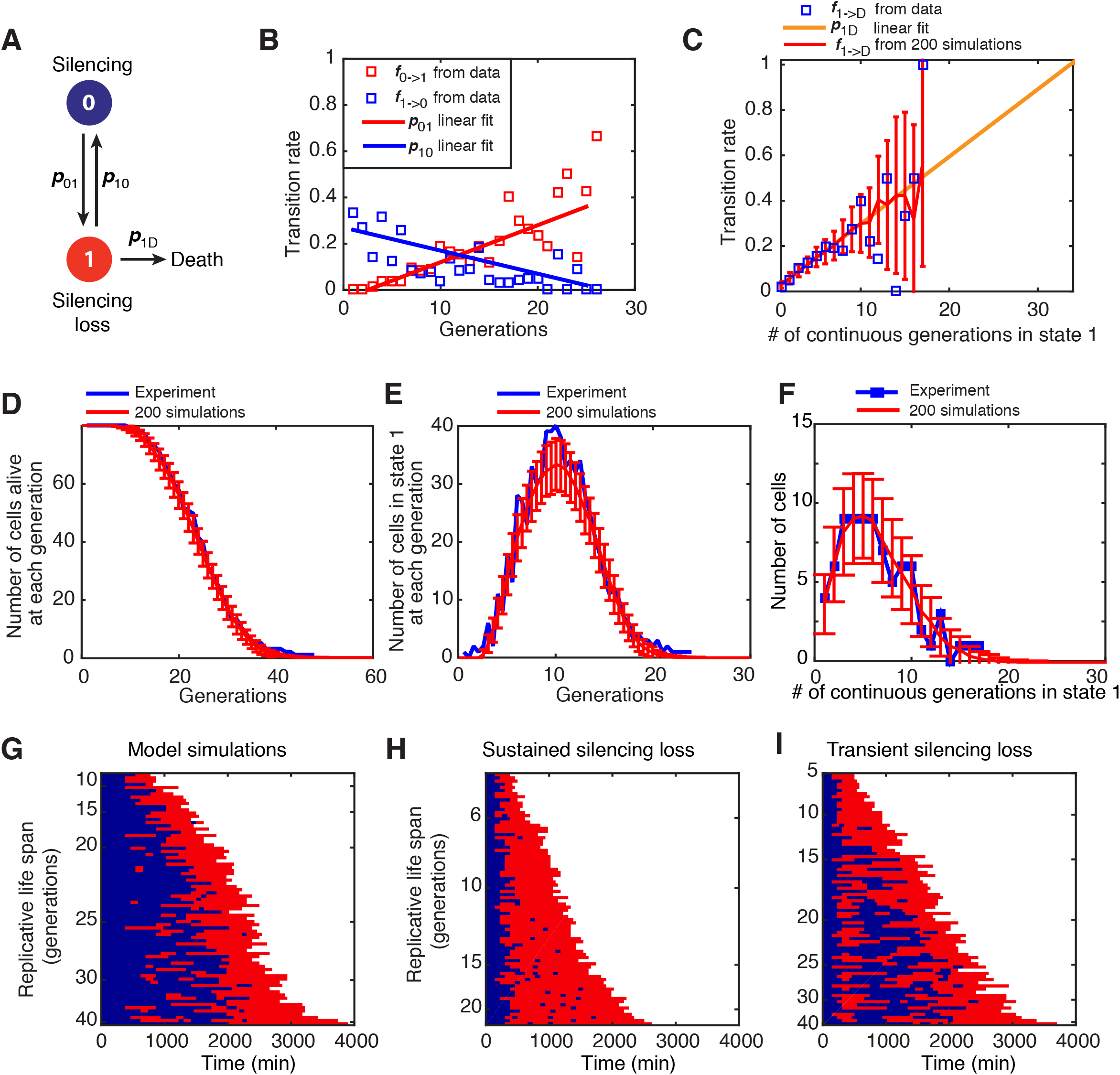
A phenomenological model of cell aging. (A) Schematic diagram of the model. The circles indicate the cellular states, and the arrows depict transitions between the states. (B) The statistics of silencing state transitions as a function of age. The fraction of all cells at state 0 of a given generation that switch to state 1 at the next cell cycle (red) and the fraction of the cells at state 1 that return to state 0 at the next cell cycle (blue) have been computed as a function of age. (C) The transition rates from state 1 to death as a function of the number of consecutive generations in state 1. Blue squares are experimentally measured fractions of cells that died exactly after *N* consecutive generation in state 1 over the total number of cells that lived for at least *N* generations. Yellow straight line is a linear fit of these data (0<*N*<10). The red line and the error bars indicate the mean and standard deviation (std) of the fraction 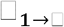 from simulations. (D) The average number of cells alive as a function of age and the std in simulations (red line and error bars) and experimental data (blue line). (E) The average number of cells in state 1 as a function of the age and the std in simulations (red line and error bars) and experimental data (blue line). (F) The distribution of the number of consecutive generations in state 1 until death in simulations (red line and error bars) and experimental data (blue line). All simulation results in (C-F) were obtained from 200 stochastic simulations of 79 cells. (G) Singlecell state trajectories from a single stochastic simulation of the model with 79 cells at time 0. Each row represents the time trace of a single cell throughout its lifespan. Blue corresponds to state 0, and red to state 1. Cells are sorted based on their RLSs. (H) Single-cell state trajectories simulated using modified transition rates to reflect sustained silencing loss. (I) Single-cell state trajectories simulated using modified transition rates for 4 generations to reflect transient silencing loss. Details of the model are included in SI Appendix.

Chromatin silencing at the rDNA is primarily mediated by the lysine deacetylase Sir2, encoded by the best-studied longevity gene to date, which is conserved from bacteria to humans (18). To examine the role of Sir2 in regulating silencing dynamics, we monitored the aging process of *sir2Δ* cells. We observed that *sir2Δ* cells do not exhibit sporadic silencing loss; instead, most cells show sustained silencing loss throughout their lifespans (Fig. 5A), indicating that the intermittent silencing dynamics is dependent on Sir2-mediated silencing reestablishment. Most (~70%) *sir2Δ* cells continuously produce elongated daughters until their death, in accordance with the observed correlation between silencing loss and elongated daughters. Furthermore, in *sir2Δ* cells, sustained silencing loss leads to cell death within a relatively uniform time frame, strikingly resembling the Sustained Phase in WT cells (Fig. 5A, red dashed line). These results suggest that Sir2 promotes longevity by generating intermittent silencing dynamics and delaying entry into the Sustained Phase. We also examined *sgf73Δ*, a mutant with an extended longevity (28), and observed intermittent silencing dynamics during aging and elongated daughters at the late phase of aging. This long-lived mutant shows more silencing loss pulses than WT, consistent with the possibility that the intermittent silencing dynamics promotes longevity (SI Appendix, Fig. S5).

**Figure 5.**
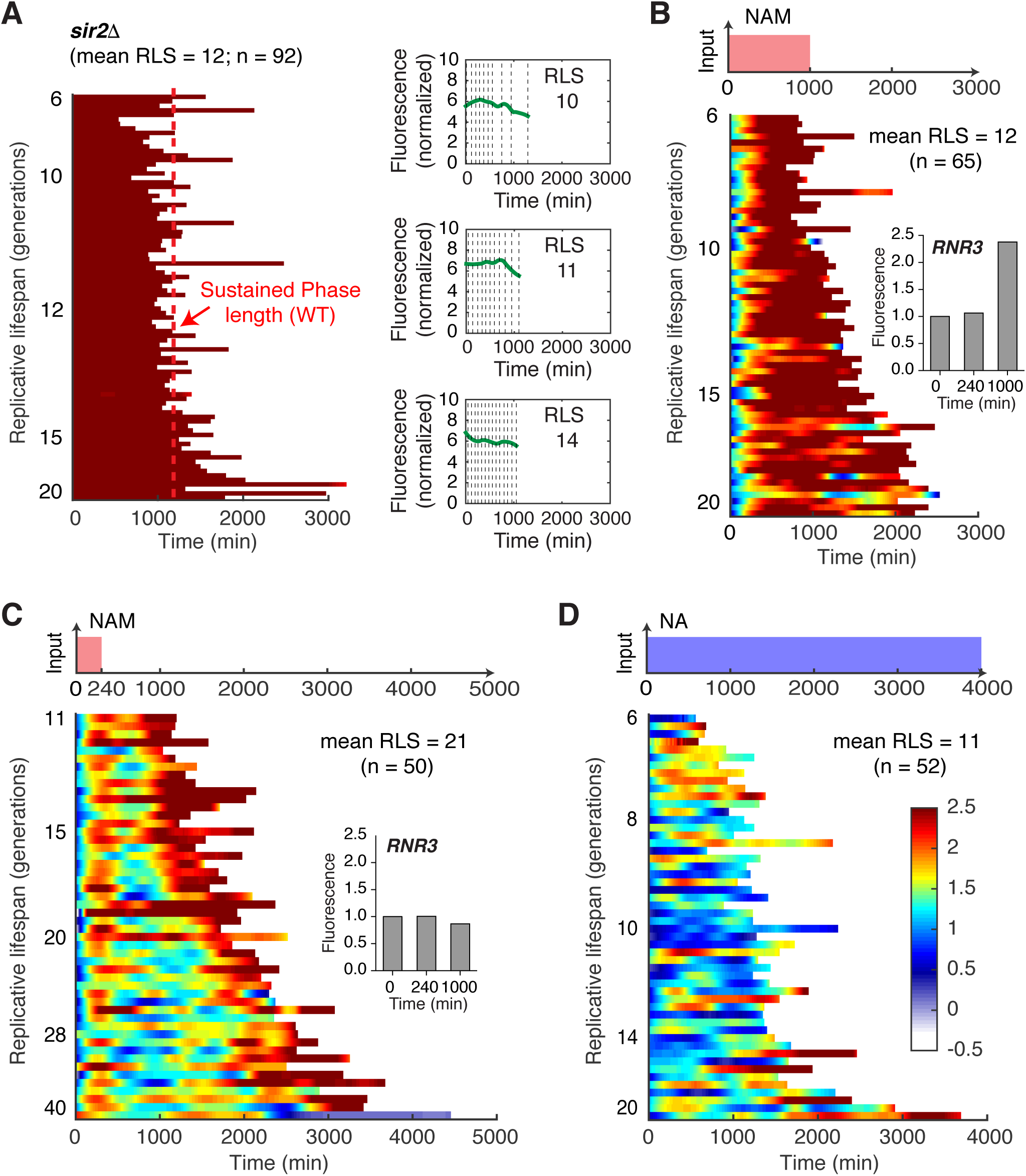
Perturbations of silencing dynamics shorten lifespan. (A) Dynamics of silencing loss in *sir2Δ* cells. Left: single-cell color map trajectories of reporter fluorescence. Each row represents the time trace of a single cell throughout its lifespan. Color bar is identical to Fig. 2. Cells are sorted based on their RLS. The red dashed line represents the average length of Sustained Phase in WT cells, defined in Fig. 3b. Right: representative single-cell time traces with different lifespans. Dynamics of silencing loss in WT cells are shown in response to chemical perturbations: (B) A 1000-min 5 mM NAM pulse. Inset shows the expression of *RNR3*-mCherry in response to a 1000-min 5 mM NAM pulse at indicated times; (C) A 240-min 5 mM NAM pulse. Inset shows the expression of *RNR3*-mCherry in response to a 240-min 5 mM NAM pulse at indicated times. (D) A constant treatment of 2.5 mM NA. For each condition, (top) a schematic illustrating the time-based drug treatment has been shown with (bottom) the corresponding single-cell color map trajectories of reporter fluorescence.

To further test predictions of the model and examine the causative roles of silencing dynamics on aging, we exposed cells to nicotinamide (NAM), an inhibitor of Sir2 (29), to chemically disrupt silencing with physiologically relevant durations. In response to a 1000-minute NAM input mimicking the Sustained Phase, the majority of cells cannot recover from silencing loss (Fig. 5B). All (100%) of the treated cells, although young, continuously produce elongated daughters and die within a similar time frame to sustained silencing loss in *sir2* mutant or WT cells (Fig. 5B). The cells also show an elevated DNA damage response, as reported by the induction of *RNR3* (30, 31) (Fig. 5B, inset), and have a significantly shortened lifespan (mean RLS: 12), comparable to that of *sir2* mutants. These results suggested that sustained silencing loss causes the elongated daughter phenotype and accelerates cell death in young cells. In contrast, in response to a 240-minute NAM input, mimicking the sporadic silencing loss, most cells exhibit a synchronized silencing loss followed by effective silencing re-establishment upon the removal of NAM (Fig. 5C). This short-term silencing loss does not induce the DNA damage response (Fig. 5C, inset) and does not affect lifespan, in accord with the sporadic silencing loss in Intermittent Phase of naturally aging cells. These perturbation experiments validate the model’s predictions and confirm that a prolonged duration of silencing loss triggers an irreversible process sufficient to induce cell death, whereas short-term silencing loss is fully reversible and does not promote aging or death.

Finally, we considered the role of the intermittent pattern of silencing, as opposed to continuous chromatin silencing. To this end, we used nicotinic acid (NA), an activator of Sir2 (32), to prevent sporadic silencing loss in aging cells. As shown in Fig. 5D, most cells show continuous silencing with few sporadic losses of silencing. The repression of silencing loss during aging results in the absence of the elongated daughter phenotype in the majority (~98%) of cells. Importantly, this sustained chromatin silencing also results in significantly shortened lifespan (mean RLS: 11), suggesting that constant chromatin silencing can activate a different aging or death pathway. Furthermore, we confirmed that the effects of NA on lifespan are not likely due to direct cellular toxicity (SI Appendix, Fig. S6), but are instead primarily mediated through Sir2 (SI Appendix, Fig. S7). Our results suggest the intriguing possibility that chromatin silencing may be a double-edged sword with both anti-aging and pro-aging functions, probably depending on target genomic regions. Whereas sustained rDNA silencing loss increases heterochromatin instability, sustained chromatin silencing may repress genes that benefit longevity; either state would accelerate cell aging and death. The intermittent silencing dynamics driven by Sir2 allow the cell to periodically alternate between the two states, avoiding a prolonged duration in either state and maintaining a time-based balance important for longevity.

## Discussion

Further analysis will continue to identify downstream targets that mediate the negative effects of sustained silencing on lifespan. For example, potential candidates include genes in other heterochromatic regions, such as subtelomeric genes that encode metabolic enzymes or have mitochondrial functions (33), or Sir2-repressed genes with pro-longevity functions (34), or critical processes influenced by rDNA transcription, such as ribosomal biogenesis, a potential regulator of yeast aging (35, 36). Interestingly, a recent study (37) demonstrated that aggregation of a cell-cycle regulator, but not the previously reported loss of silencing at *HM* loci (38), causes sterility in old yeast cells. This work, together with our findings here, suggests that chromatin silencing at various genomic regions might undergo different age-dependent changes, probably due to their specific silencing complexes. For example, whereas the silencing at *HM* loci is regulated by a protein complex containing Sir2, Sir3 and Sir4 (39), a different complex containing Sir2, Net1, Cdc14, and Nan1 is required for the silencing at the rDNA (40, 41). Furthermore, it has been shown that rDNA and *HM* silencing have different sensitivities to NAM or genetic perturbations (42–44), implying different regulatory modes at these loci. We anticipate that further systems-level analysis will enable a more comprehensive understanding of chromatin regulation during aging. Another interesting question for future investigation is the fate of elongated daughters. One possibility is that the production of these abnormal daughters may serve as a rescue strategy to alleviate damage accumulation in mother cells. Future technological advances that allow lifespan tracking of selected daughter cells would enable us to examine the silencing dynamics, lifespan and aging type decision of these elongated daughters and their daughters.

Dynamics-based regulation is an emerging theme in biology, the role of which has been increasingly appreciated in many biological processes across a wide range of organisms (3, 4, 45–49). Our analysis here uncovered the significant role of silencing dynamics in cellular aging and opened the possibility of designing temporally controlled perturbations to extend lifespan. For example, although constant NA exposure shortens lifespan, it might be possible to design dynamic regimes of NA treatment to specifically prevent sustained silencing loss and thereby delay aging. Given the observed single-cell heterogeneity in silencing and aging dynamics (Figs. 2 and 3), future efforts will focus on new technologies that enable distinct real-time NA treatments to individual cells based on their silencing states.

## Materials and Methods

### Strains and plasmids construction

Standard methods for the growth, maintenance and transformation of yeast and bacteria and for manipulation of DNA were used throughout. The yeast strains used in this study were generated from the BY4741 (*MAT a his3Δ1 leu2Δ0 metî5Δ0 ura3Δ0*) strain background. Strain and plasmid information is provided in Tables S1 and S2.

### Microfluidic device for yeast aging studies

In designing a microfluidic device for studying aging in budding yeast, the viability of the cells, efficiency of cell trapping and robustness of the device were our primary concerns. The robustness of the device is affected by clogging due to excess cells around the traps and at the waste port, which can interfere with mother cell lifespan and retention. Supplying media through ~20 μm tall main channels readily allowed excess cells to be washed away and prevented clogging. Therefore, a critical feature of our device is its two-layer design, making it extremely robust over the course of our experiments that takes more than 80 hours. This is a unique feature compared to recently published devices that are all single-layer (10, 13, 14). The device was optimized for using continuous gravity-driven flow during operation, with the three-inlet design also facilitating media switching experiments. Cell loading efficiencies and final retentions until cell death are approximately 93% and 75% respectively. See SI Appendix for further details on the device and its validation.

### Live-cell imaging and analysis

Time-lapse imaging experiments were performed using a Nikon Ti-E inverted fluorescence microscope with Perfect Focus, coupled with an EMCCD camera (Andor iXon X3 DU897). The light source is a spectra X LED system. Images were taken using a CFI plan Apochromat Lambda DM 60X oil immersion objective (NA 1.40 WD 0.13MM). The microfluidic device was placed on the motorized microscope stage (with Encoders) within a 30°C incubator system. The flow of medium in the device was maintained by gravity and drove cells into traps. Waste medium was collected to measure flow rate, which was about 2.5 ml/day. Images were acquired every 15 min for a total of 80 or more hours. Images were processed and quantified with a custom MATLAB code. Cell divisions of each mother cell were manually identified and counted at the time that the nuclei separated between mother and daughter cells. Cells were categorized based on their aging phenotypes, characterized by the morphologies of later daughters they produced. Mothers continually producing elongated daughters at the last few generations were categorized as “aging with elongated daughters”, whereas mothers continually producing round daughters at the last few generations were categorized as “aging with rounded daughters.” Dynamic patterns of reporter fluorescence have been shown in ‘jet’ rainbow color maps. The ‘inferno’ versions of the color maps to provide continuous luminance visualization of the data have been included in SI Appendix, Fig. S8.

Detailed methods and the development of the phenomenological model are given in SI Appendix.

## Acknowledgments

We thank Yuan Zhao (Bioinformatics, UCSD) for helping with developing the image analysis code, and Dr. Philbert Tsai (Physics, UCSD) for helping with the superresolution confocal microscopy. This work was supported by NSF MCB-1616127 (to N.H.), UC Cancer Research Coordinating Committee (to L.P.) and DoD, Air Force Office of Scientific Research, National Defense Science and Engineering Graduate (NDSEG) Fellowship, 32 CFR 168a (to R.O.).

## Author Contributions

Y.L. conducted the experimental analysis. M.J. and P.B. developed the computational model. R.O. developed the microfluidic device. Y.L., M.J., and R.O. performed data analysis. Y.L., M.J., R.O., L.S.T, L.P., J.H. and N.H. conceived the project and wrote the paper.

